# Large polyacrylamide hydrogels for large-batch cell culture and mechanobiological studies

**DOI:** 10.1101/2023.01.27.525967

**Authors:** Xuechen Shi, Paul Janmey

**Affiliations:** Institute for Medicine and Engineering and Department of Physiology, University of Pennsylvania, PA 19104, USA

## Abstract

The rigidity of a cell’s substrate or extracellular matrix plays a vital role in regulating cell and tissue functions. Polyacrylamide (PAAm) hydrogels are one of the most widely used cell culture substrates that provide a physiologically relevant range of stiffness. However, it is still arduous and time-consuming to prepare PAAm substrates in large batches for high-yield or multi-scale cell cultures. Here we present a simple method to prepare PAAm hydrogels with less time cost and easily accessible materials. The hydrogel is mechanically uniform and supports cell culture in a large batch. We further show that the stiffness of the hydrogel covers a large range of Young’s modulus and is sensed by cells, regulating various cell features including changes in cell morphology, proliferation, and contractility. This method improves reproducibility of mechanobiology studies and can be easily applied for mechanobiology research requiring large numbers of cells or experimental groups.

## Introductions

Polyacrylamide (PAAm) hydrogel substrates have been broadly used as a two-dimensional mammalian cell culture system. Compared with other natural or synthetic polymeric materials, PAAm has several unique characteristics: (1) tunable elasticity or viscoelasticity ^[1]^; (2) resistance to degradation, providing stable mechanical properties over time; (3) controllable and constant surface biochemistry ^[2]^; and (4) translucence and low auto-fluorescence. These features make PAAm hydrogels a good platform to investigate how cells and tissues respond to extracellular matrix rigidity as well as to measure cellular force generation and transmission. Examples of the biological processes and/or properties studied include cell adhesion and morphology ^[3]^, durotaxis ^[4]^, traction force ^[5]^, cell differentiation ^[6]^, wound healing ^[7]^, endocytosis ^[8]^, plithotaxis ^[9]^, cancer metastasis ^[10]^, elastocapillarity ^[11]^, etc. All these studies suggest a role for the PAAm gel system in fundamental mechanobiological research as well as for potential applications such as drug screening ^[12]^, tissue engineering ^[13]^, and cancer therapies ^[14]^.

In preparation of the PAAm substrate, a PAAm hydrogel is typically polymerized between an adhesive glass slide and a non-adhesive hydrophobic or silanized glass coverslip. The polymerized hydrogel is thus physically or covalently attached to a rigid substrate at its bottom surface, and the hydrophobic slide is removed to expose the cell-contacting surface. However, this approach as well as other modified methods ^[13a, 14a, 15]^ is associated with difficulties, especially for cell culture in large batches. For example, the glass-sandwich method is technically demanding requiring several chemical reactions that take multiple experimental steps. The mechanical imbalance between an adhesive surface on the bottom and a free, fluid-exposed surface on the top can cause wrinkling or gel damage if the gel swells in culture medium. An erroneous handling in any step could result in the gel peeling off from the coverslip or roughening of the upper surface in later experimental stages. Such time-consuming procedures also demand efforts to prepare substrates in large numbers, making it very challenging to conduct biochemical or proteomic studies that require million-level cells. Finally, as the pre-gel solution for each piece of PAAm gel is typically in a small volume, e.g. 25 μl, mechanical variations may arise since the diffusion of oxygen profoundly inhibits acrylamide polymerization and exposure to air is difficult to regulate uniformly.

Here we present a novel method to prepare a PAAm hydrogel as a cell culture substrate to overcome the above issues. This one-simple-step method allows the production of a PAAm hydrogel with a large surface area, using immediately accessible materials and taking a short period of time. Upon swelling, the hydrogel can be cut into custom sizes for various purposes. We show that the PAAm hydrogel synthesized by this technique is mechanically uniform and experimentally reproducible. The surface of the hydrogel can be functionalized with cell adhesion ligands so that cells may grow to form a monolayer on this large hydrogel, generating milligram-levels of total proteins. Finally, we show that the stiffness of the hydrogel can be perceived by cells, resulting in various cell responses. The novel technique presented here offers an adequate platform to support research in mechanobiology and may serve as an alternative method to the traditional glass-sandwich protocol.

## Results

### One-step synthesis of large PAAm hydrogel

Our method prepares the large PAAm hydrogel between two polystyrene surfaces with a 100 mm, large polystyrene petri dish. As illustrated in **Fig. 1A**, the pre-gel solution containing acrylamide, bis-acrylamide, APS, and TEMED is transferred into a plastic petri dish lid with the opening side facing up. The bottom half of the petri dish, likewise opening facing up, is then placed onto the pre-gel solution covering it. In this way, the hydrogel is sandwiched between the two petri dish surfaces (**Fig. 1B**). Because the size of the petri dish lid is slightly larger than its body, the boundary of the synthesized hydrogel is exposed to air, which inhibits acrylamide polymerization, and is thereafter discarded in the following steps. We use higher concentrations of the initiator APS and catalyst TEMED compared with the commonly used formulation ^[16]^ to minimize potential ambient mechanical perturbations, as the dish body is free-floating on the pre-gel solution before gelation happens.

**Figure 1.**
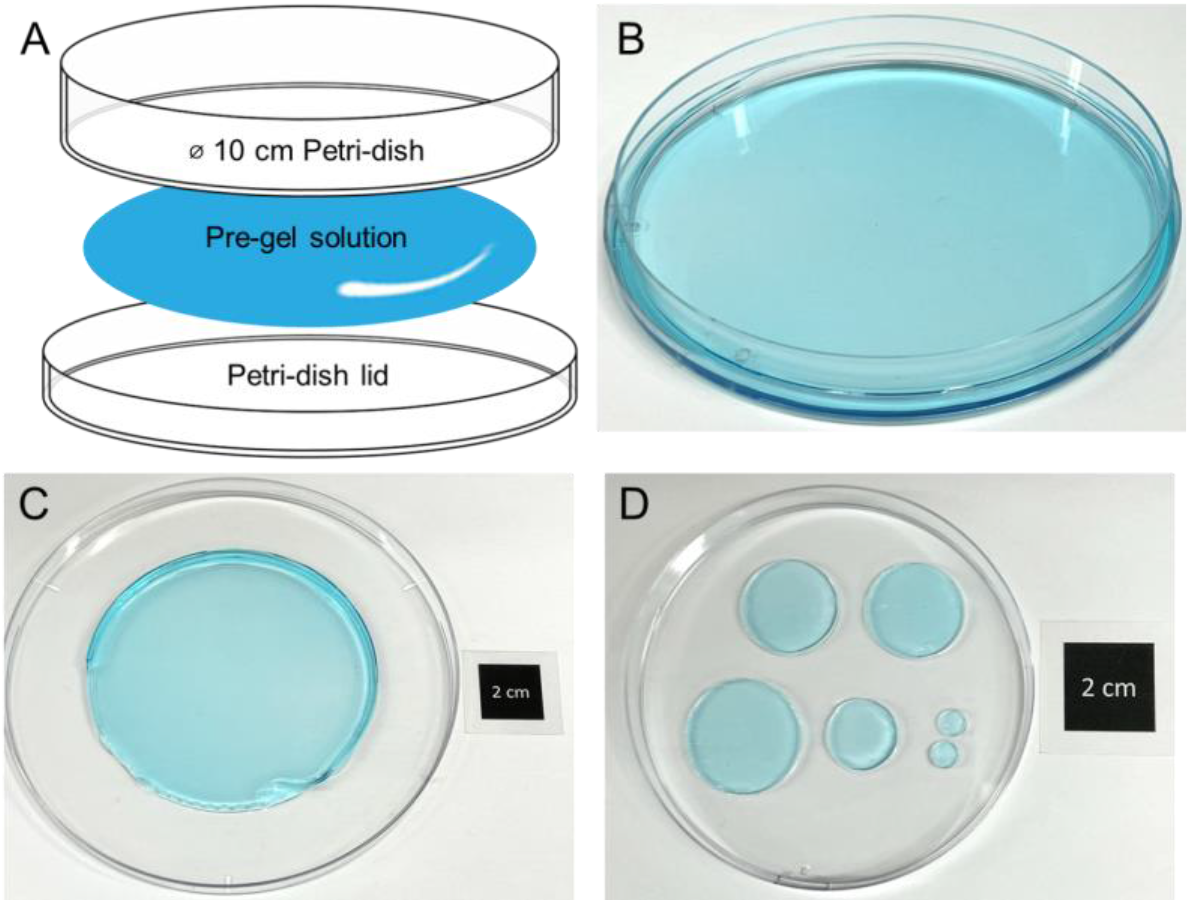
Synthesis of a large PAAm hydrogel with the plastic petri dish. **A**. Schematic view of the one-step synthesis method. The pre-gel solution is sandwiched between two surfaces of a large petri dish and its lid. **B**. Photograph of the hydrogel synthesizing setup. A droplet of blue food color is mixed in the pre-gel solution for the purpose of illustration only. **C**. Photograph of the synthesized large hydrogel after taking it from the dish-lid setup. The gel, mixed with blue food color, is placed in a larger container during photography. **D**. Photograph of small pieces of gels cut from the large hydrogel with tissue punchers of different sizes.

After polymerization in a humidity chamber for 30 minutes, the top dish is removed, and the hydrogel is taken out from the dish lid. A splash of water may help this peeling-off process. The resulting hydrogel, except for the out-of-dish boundary region, is transparent, flat with no curling. The hydrogel can be used in its large dimension (**Fig. 1C**), or can be cut into multiple pieces with different sizes per future experiments needs (**Fig. 1D**). Note that to acquire a desirable size, the cutting should be done based on the swelling ratio or after the hydrogel fully swells.

The non-treated petri dish is commercially available and inexpensive. For this petri dish with nominal 100 mm diameter, the diameters of its body and lid are 88 mm and 94 mm, respectively. Unlike the glass-sandwiching method which requires hydrophobic and gel-attaching treatments to the glass slides, our method does not involve any surface treatment, as the petri dish is non-plasma-treated and is hydrophobic. Together, our method is cost-effective yet generating a relatively large-sized hydrogel in the meantime.

### Mechanical properties of large PAAm hydrogels

To determine the mechanical properties of the large PAAm substrate synthesized by our novel method, here we prepare hydrogels with three stiffnesses by varying the acrylamide and bis-acrylamide concentrations with the formations in **Table 1**. The total volume of pre-gel solution for each gel is 12 ml, and the resulting hydrogel thickness is around 1.5 mm, although softer gels are slightly thicker due to swelling. After being taken out from the dish lid, the hydrogels are with no constraint, thus they swell when immersed in an aqueous environment. This changes their sizes and mechanical properties ^[17]^. For cells to receive constant mechanical input, it is necessary to pre-swell the hydrogels until they reach their equilibrium swelling states. To address this, we first characterize their swelling dynamics. The swelling test was done in PBS at 37°C, providing gels with the same osmotic pressure and temperature as those during cell culture. We find that the softer gels take a longer time to reach equilibrium compared to the stiffer gels, and the softest gel takes around 24 hours to swell to its final volume **(Fig. 2A)**. With these results, in the following experiments we wash and incubate the hydrogels for at least 36 hours before the next steps.

**Table 1.**
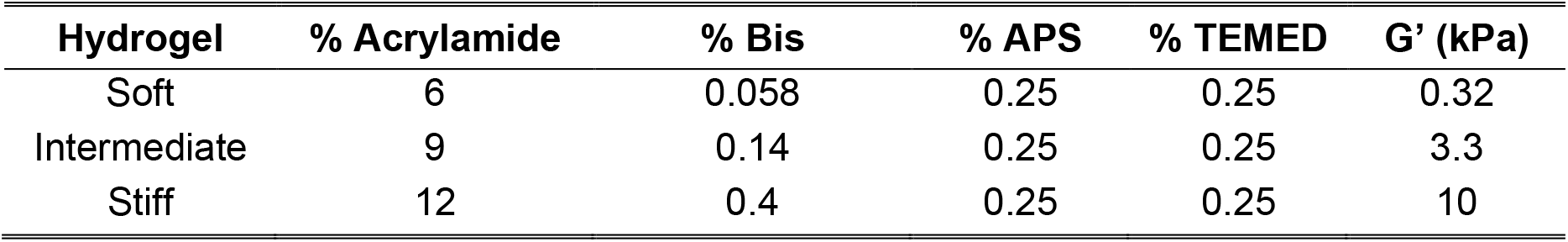
Formulations for synthesis of large polyacrylamide hydrogels and their expected elastic shear moduli G’.

**Figure 2.**
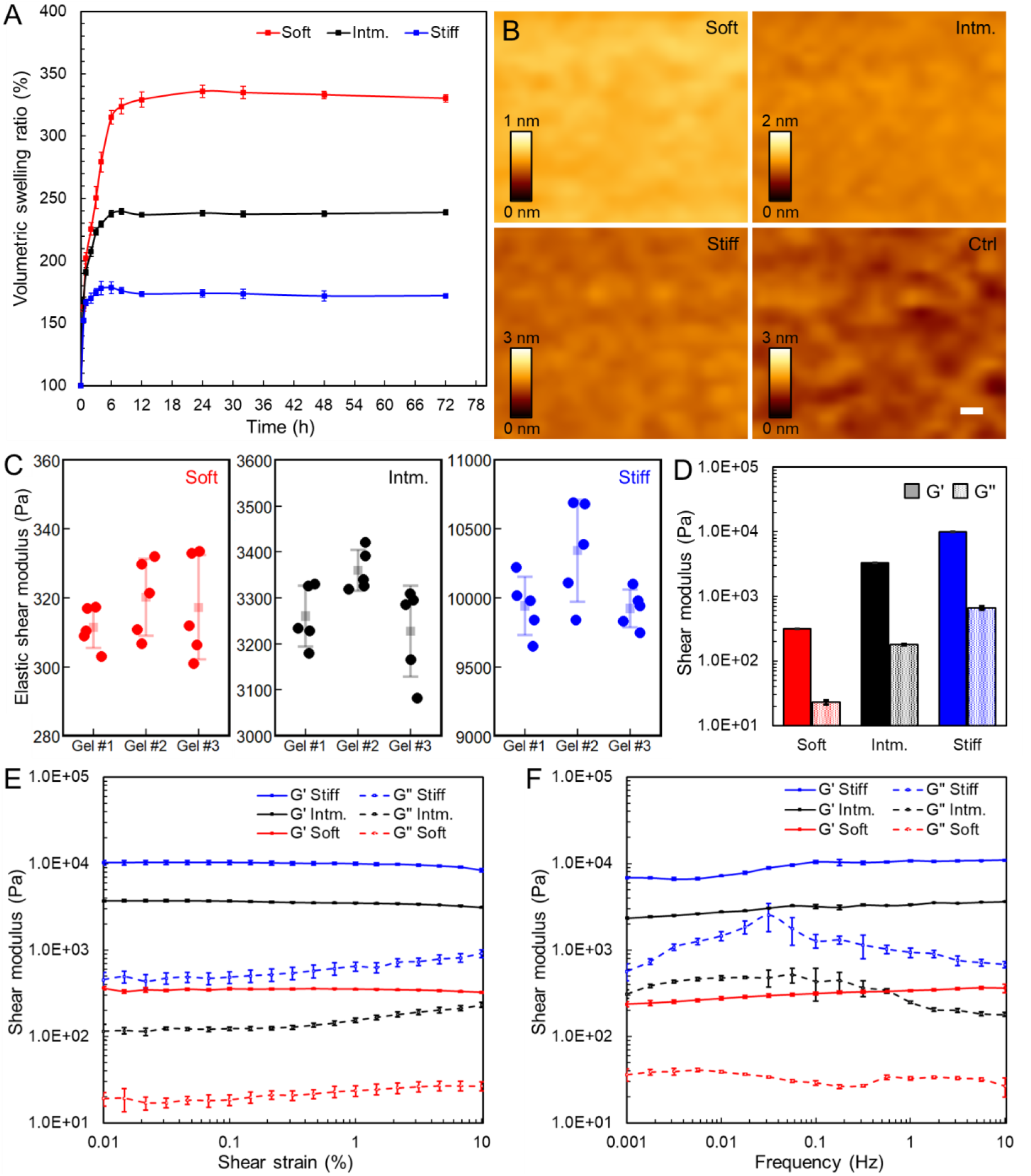
Mechanical characterization of large PAAm hydrogels. **A**. Dynamics of the volumetric swelling ratio of the hydrogels. Hydrogels swell in PBS at 37°C. For each stiffness, n=6 from three independent large hydrogels. Error bar denotes standard error of the mean (SEM). **B**. AFM images for the surface of the large hydrogels. Ctrl: control image from the hydrogel polymerized between glass slides. Scale bar=50 nm. **C**. Elastic shear moduli of the hydrogel pieces randomly taken from the large hydrogels. For each stiffness, three independent large hydrogels are prepared, and n=5 circular pieces with 20 mm diameter punched from each large hydrogel are tested. Each solid dot denotes the measured elastic shear modulus of each gel piece; light-colored square denotes the average value and error bar denotes standard deviation.**D**. Average shear moduli of the gel pieces taken from large hydrogels. Solid and dashed columns denote the elastic (G’) and viscous (G”) shear moduli, respectively. For each stiffness, n=15 from three independent large hydrogel. Error bar denotes SEM. Tests in C and D are conducted with 1% strain at 1 Hz. **E** and **F**. Average shear modulus of the hydrogel pieces as a function of the shear strain and oscillation frequency, respectively. Solid and dashed curves denote the elastic (G’) and viscous (G’’) shear moduli, respectively. For each stiffness, n=3 from three independent large hydrogel. Error bar denotes SEM.

We next demonstrate that the surface roughness of the large PAAm gels is small, similar to that of the traditional PAAm hydrogel polymerized between glass slides, as imaged by atomic force microscopy (AFM, **Fig. 2B**). To determine the mechanical uniformity of the large PAAm gels, we prepared a total of three large gels for each stiffness, and across each large gel we cut five hydrogel pieces at random locations with a circular 20 mm-diameter tissue puncher. The hydrogel stiffness was measured by a rheometer with 1% shear strain at 1 Hz at 37°C. We find that for each stiffness of formulation, there is less than 5% variance in their elastic shear moduli G’ across one large gel and less than 2.5% variance among three parallel gels **(Fig. 2C)**. We therefore conclude that the hydrogel is mechanically uniform and experimentally reproducible. The combined rheology results **(Fig. 2D)** show that the Young’s modulus of the gels ranges from 1 to 30 kPa, covering the majority of physiologically appropriate stiffnesses ^[2]^. The softest we have made has a shear modulus of 266 Pa; while in theory it’s possible to further decrease the stiffness by adjusting the formulations, softer gels may become difficult to handle and transfer. In addition, we note that there is an approximately 6% loss tangent in our hydrogels **(Fig. 2D)**, compared to <1% for PAAm gels formed directly between rheometer plates, the reason for which is unclear to us. The amplitude sweep test indicates that the hydrogel stiffness is almost independent of the applied strain **(Fig. 2E)**. The frequency sweep test indicates that the hydrogel stiffness increases very slightly with the applied frequency **(Fig. 2F)**. The shear modulus increases by around 20% with a 100× change in frequency ranging from 0.1 to 10 rad/s for the soft and intermediate gels, and around 35% for the stiff gel, where 1 rad/s=0.159 Hz is generally regarded as a biologically relevant frequency ^[1]^.

For all experiments across this manuscript, we used the fully swollen, free-floating hydrogel. On the other hand, the hydrogel can also be used inside the dish lid without taking it out from the lid upon gelation. For gels held within the petri dish, the hydrogel does not undergo x-y swelling, thus the formulation for a specific rigidity is different from the formulation of a boundary-free hydrogel. Secondly, as the hydrogel boundary is exposed to air during polymerization, experiments within that region should be avoided. Thirdly, during cell culture, a larger container may be needed to hold the lid-containing hydrogel and to cover the cell culture medium, which cannot be simply achieved by the same petri dish.

### Protein functionalization at the hydrogel surface

We further study whether the large hydrogel can be functionalized with cell adhesive ligands to enable cell adhesion. To this end, we first activated the hydrogel surface with the bi-functional crosslinker Sulfo-SANPAH. The fully swollen gel was tapped dry and, with its size obviously larger than its original 100 mm in diameter due to swelling, punched by the 100-mm petri dish prior to the activation. Sulfo-SANPAH solution in 2 ml was spread on the hydrogel surface **(Fig. 3A)** followed by UV treatment. We next use fluorophore-labelled protein to determine the protein coating efficacy. After the gel was washed to remove the excess crosslinker, the gel was coated with rhodamine-labelled fibronectin for 2 hours. The hydrogel was washed for one day and was imaged by an upright fluorescence microscope. The fluorescence signal was detected on the hydrogel surface **(Fig. 3B)**, suggesting that like the traditional PAAm hydrogel sandwiched between glass slides, cell adhesive proteins can be attached to our large hydrogel.

**Figure 3.**
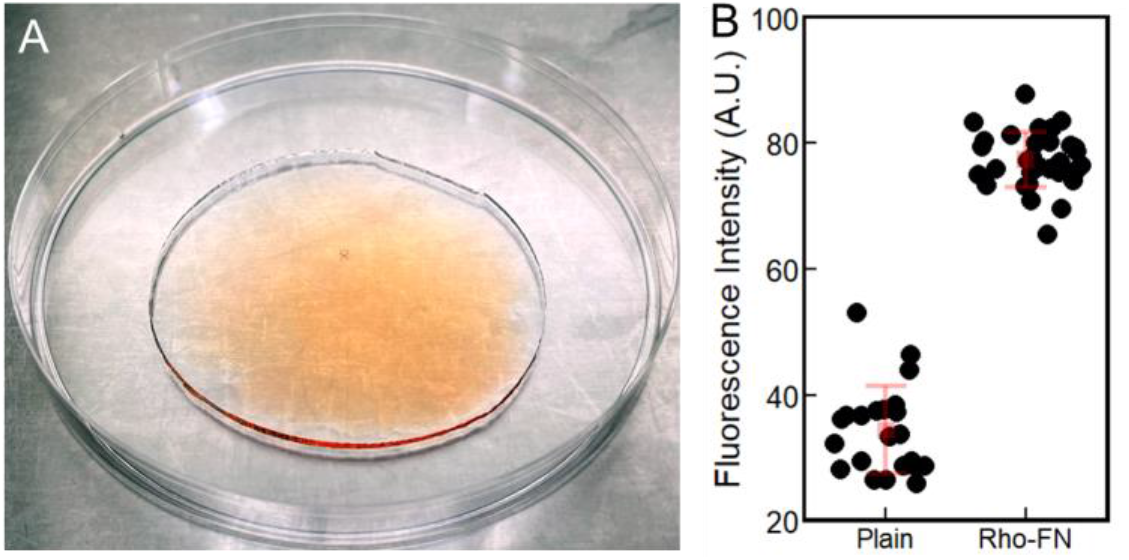
Protein attachment to the large hydrogel surface. **A**. Photograph of spreading Sulfo-SANPAH solution onto the large hydrogel for protein crosslinking. **B**. Fluorescence intensity of the large hydrogel after crosslinking with rhodamine-labelled fibronectin. Fluorescence images are taken at randomly chosen locations across the entire large hydrogel. The pink square denotes the average value, and the error bar denotes standard deviation.

### Large-batch cell culture on large PAAm substrate

One of the motivations for the method we present here is to generate a large cell growing area with controlled substrate rigidity yet costing a short amount of time. This is useful for research like proteomics that typically require intracellular proteins in nano- or microgram level and million-scale of cells. To assess whether cells could adhere and grow on the hydrogel substrate and whether this substrate enables cell culture in a large batch, we coat the large hydrogel with collagen I and seed bovine aortic endothelial cells (BAEC) on top of the gel. Cells were cultured for three days at which time they became confluent **(Fig. 4A)**, forming a monolayer with nematic, parallel array-like structures locally ^[18]^. Cells were then trypsinized, washed, and lysed. The total cell number and total cellular protein amount harvested was determined to be 9.4 million and 0.94 mg, respectively. 20 μg whole cell proteins were resolved by SDS-PAGE and stained with Coomassie blue **(Fig. 4B)**. To further demonstrate that the large-batch cell culture on the large PAAm substrates could produce ample amount of whole cell proteins for protein analyses, we conduct Western blot experiment especially targeting on the low-abundance proteins. Human mesenchymal stem cells (hMSC) were cultured on collagen coated PAAm substrates with various rigidities or in a polystyrene petri dish for three days **(Fig. 4C)**. The total cellular protein amount from each substrate was: 0.18 mg for Soft, 0.26 mg for Intermediate, 0.30 mg for Stiff, and 0.25 mg for PS. Cell lysate from each substrate was normalized by the total protein concentration, and 17 μg of whole cell proteins were loaded for electrophoresis. The lower-abundance proteins, including filamin A, gelsolin, and lamin A and C, were visualized in Western blotting together with the highly abundant proteins such as actin and vimentin **(Fig. 4D)**, and yet this only took a small portion of the whole cell proteins harvested. The advantage of detecting multiple proteins from single preparations of cell extracts is illustrated by the significantly lower amount of full length filamin A in the extract prepared from cells on polystyrene, compared to the amounts on collagen-coated gels. The similar or larger amounts of other proteins such as actin or vimentin serve as controls for possible differences in total protein. The decreased amount of filamin A on polystyrene is possibly related to the binding of cells on plastic primarily through fibronectin or vitronectin receptors that engage primarily talin, rather than collagen receptors that engage primarily filamin ^[19]^, leading either to decreased expression or increased degradation of the protein. Taken together, these results show that our large hydrogel supports large-scale and high-yield cell culture for protein studies

**Figure 4.**
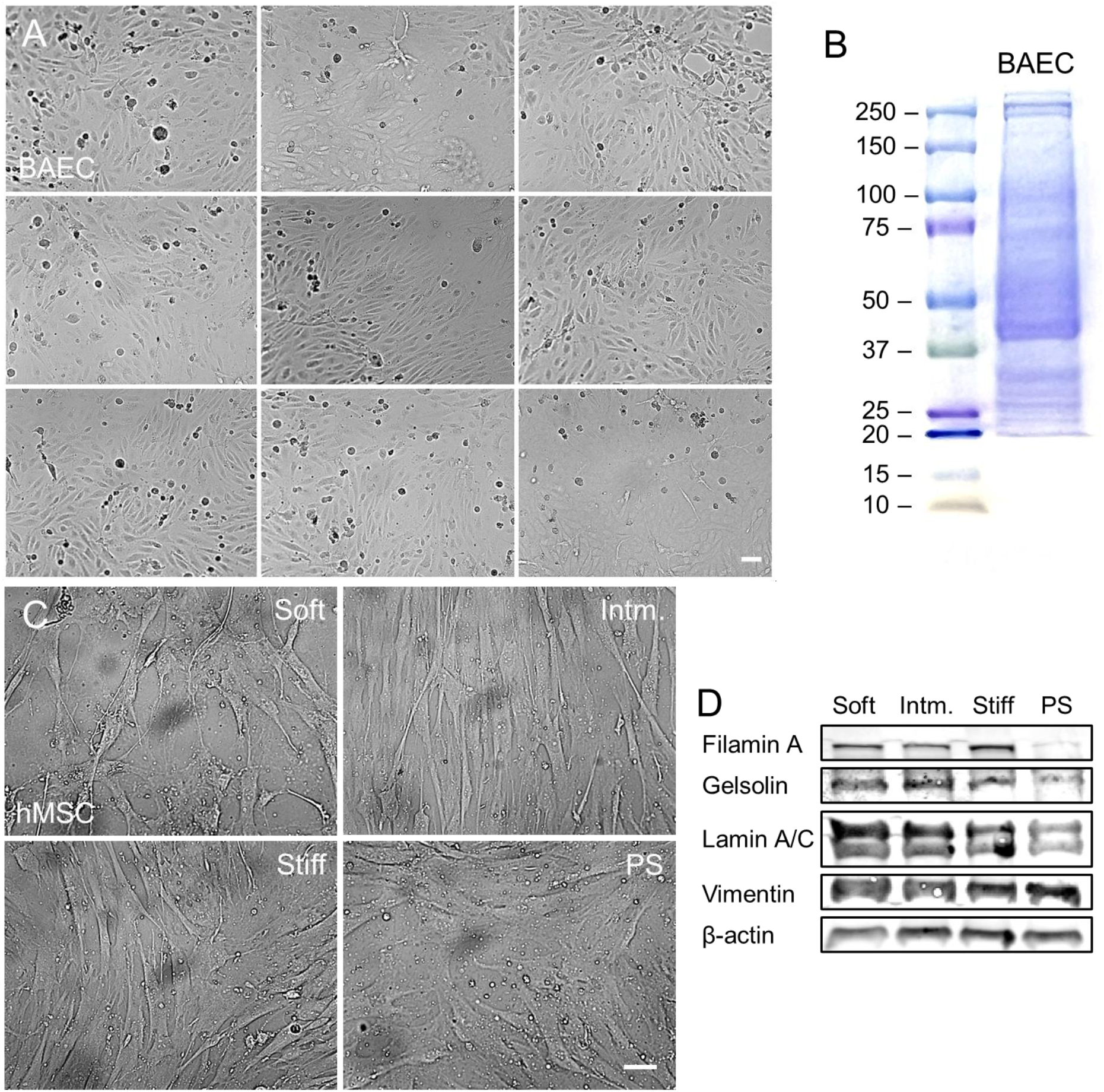
Cell culture in large batches on large PAAm substrates. **A**. Bright field images of BAEC monolayer cultured on the large hydrogel. Nine representative images are taken at randomly chosen locations across the large hydrogel. The hydrogel was prepared with the intermediate stiffness and functionalized with collagen I. **B**. Whole-cell proteins SDS-PAGE image stained with Coomassie blue. **C**. Representative bright field images of hMSCs cultured on large PAAm gels with different rigidities or polystyrene (PS) substrate for 3 days. For A and C, scale bar=50 μm. **D**. Western blot analysis of hMSCs protein expressions regulated by substrate rigidities. The total protein loading amount was consistent for each lane.

### PAAm gels as a mechanobiological research platform

We further ask whether the PAAm hydrogel synthesized by our novel method could be applied as a platform for mechanobiological research. To assess this feature, we study the cells’ responses to hydrogels with different stiffnesses. For the following cell experiments, we used circular gel pieces of 20 mm diameter punched from fully swollen large hydrogels, and we used the mechano-sensitive mouse embryo fibroblast (mEF) cell line as our cell model. Gel pieces were functionalized with collagen I prior to cell seeding. After a one-day incubation, cells on stiffer gels exhibit a significantly larger spreading area compared to cells on softer gels **(Fig. 5A** and **B)**. Fluorescent staining of F-actin also confirms the morphological difference **(Fig. 5C)**. Furthermore, cells have a higher growing rate when cultured on stiffer gels **(Fig. 5D)**. Finally, by applying traction force microscopy on the hydrogel pieces, we determine that fibroblasts are 10 times more contractile on the stiffest gels than on softest gels **(Fig. 5E** and **F)**. Given the good agreement between our results and the well-established cell mechano-responses ^[20]^, we demonstrate that our novel hydrogel preparation method can support research in biomechanics and mechanobiology.

**Figure 5.**
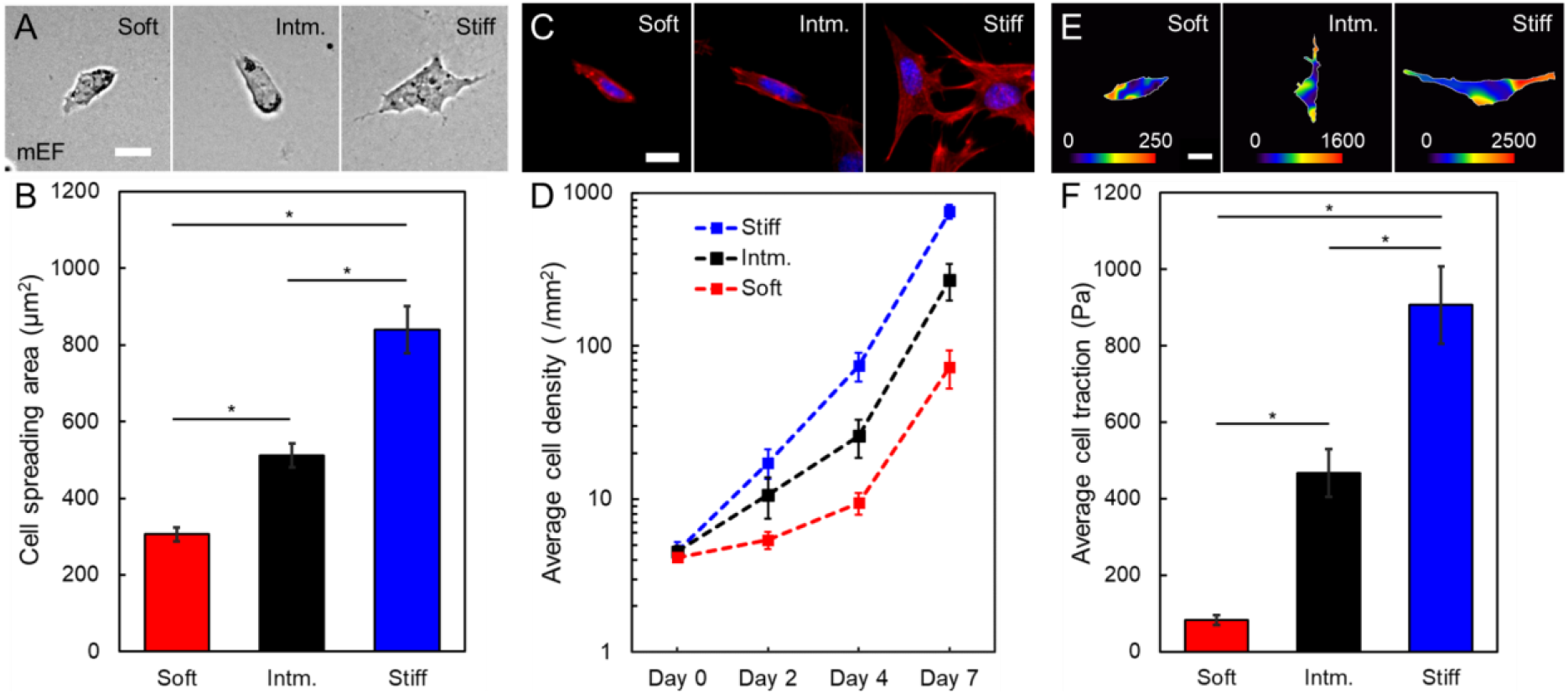
Cell mechano-responses to PAAm substrate rigidities. **A**. Representative images of mEFs after 24 h culture on gels with different rigidities. **B**. Average cell spreading area as a function of hydrogel rigidity. For each rigidity, n>40 cells on six hydrogel pieces from three independent large hydrogels. **C**. Representative fluorescence images of actin cytoskeleton in mEFs cultured on gels with different rigidities. Red: actin filament stained with phalloidin; blue: nucleus stained with DAPI. **D**. Cell proliferation rate regulated by substrate rigidity. The proliferation of mEF was determined by measuring average cell densities at different timepoints. For each rigidity, n=6 images on six hydrogel pieces from three independent large hydrogels were measured. **E**. Representative traction force maps of mEFs cultured on gels with different rigidities. **F**. Average mEF traction force as a function of hydrogel rigidity. For each rigidity, n>10 cells on six hydrogel pieces from three independent large hydrogels. For all images, scale bar=20 μm. For all graphs, error bar denotes standard error of the mean. *p<0.001 by unpaired, two-samples Student’s t-test.

## Conclusions

In conclusion, we demonstrate a new method to synthesize rigidity-tunable PAAm gel cell culture substrates. This novel method requires much less time and fewer steps to carry out and uses mechanobiological inexpensive, commonly accessible experimental supplies. The synthesized large hydrogels are not only mechanically uniform and experimentally reproducible, but also support growth of large cell populations, generating a high amount of whole cell proteins, which makes it possible to detect proteins of relatively low expression. By cutting the large hydrogel into multiple small pieces, this method also enables the mass production of substrates with different rigidities. Finally, we show that the rigidity of the hydrogels synthesized by the new method regulates various cell behaviors, which agrees with past studies using PAAm or other substrates, suggesting that the mass production method can be applied towards mechanobiology research. Altogether, we present a much easier and more cost-effective way to generate PAAm hydrogel substrates, which may be highly useful in future studies on cellular forces and other cell responses to substrate stiffness.

## Methods

### Preparation of PAAm hydrogel substrate

The hydrogel formulations are listed in **Table 1** with the volume for each large hydrogel being 12 ml. Prior to the initiation of polymerization, the pre-gel solution containing acrylamide and bis-acrylamide (Bio-Rad, Hercules, CA) is de-gassed under vacuum for 30 min. Immediately after adding APS (Fisher Scientific, Pittsburgh, PA) and TEMED (Bio Basic, Amherst, NY), the pre-gel solution is transferred onto the lid of a 100 × 15 mm petri dish (BP94S-01, Corning Inc., Corning, NY). The solution is then covered by the body of the same petri dish. This step is done with care to avoid introducing air bubbles. The entire setup is then taken into a closed chamber to maintain humidity. After 30 minutes, water of a few milliliters is added to the open edge of the hydrogel, making it easier to peel off the plastic surfaces. The gel is then taken out from the dish lid. Except for the swelling test, the hydrogels are washed in PBS for at least 36 hours at 37°C on a shaker. The washing buffer is changed for at least three times, and the total volume of the washing buffer is at least 40 times the total volume of the hydrogels. To transfer the hydrogels, we use a thin flat plastic sheet for the large gel, and a disposable plastic spoon for the 20-mm gel piece.

To attach extracellular matrix proteins, the hydrogel is first activated with sulfosuccinimidyl 6-(4′-azido-2′-nitrophenylamino) hexanoate (Sulfo-SANPAH, Pierce, Thermo Fisher Scientific, Waltham, MA). Here the remaining washing buffer around the gel edge is removed to ensure the coating solution can be held on top of the gel. 0.5 mg/ml Sulfo-SANPAH solution is spread on the gel surface in 2 ml for one large hydrogel and 0.1 ml for one 20-mm gel piece. The solution is then immediately radiated by 365 nm UV with a UV lamp for 5 minutes. After removing the excess Sulfo-SANPAH solution and washing with PBS for 40 min, the hydrogel surface is further coated with 0.1 mg/ml Rhodamine-labelled fibronectin or collagen I (Rat tail, Corning Inc., Corning, NY), again in 2 ml for one large hydrogel and 0.1 ml for one 20-mm gel piece, for 2 hours. The hydrogel is washed for one day in PBS before experiments.

### Measurement of volumetric swelling ratio

The swelling test is conducted on a shaker at 37°C with 95% humidity. Unwashed gels after polymerizing are cut into 20 mm round pieces. Gels are then immediately immersed into pre-warmed PBS when the test is started. The diameters of the hydrogel pieces are measured at different time points, during which the swelling buffers are changed. We assume the hydrogels undergo a uniform, isotropic swelling in all dimensions, and thus the unconstrained volumetric swelling ratio is calculated as J^(u)^ = V_t_ /V_0_ = (d_t_ /d_0_)^3^, where V and d are hydrogel volume and diameter, respectively, and subscripts t and 0 refer to the time point during experiment and the initial state, respectively.

### Atomic force microscopy (AFM)

A nanowizard 4 AFM mounted on a microscope was used to image the surfaces of the large PAAm hydrogels and to determine the surface roughness. The AFM was operated in contact mode with silicon nitride cantilevers (Bruker, Camarillo, CA) with a spring constant of 0.24 N/m. The height images were captured with a scan rate of 1 Hz and were processed with the build-in Pixel Difference filter to generate the surface images.

### Measurement of hydrogel rheology

Hydrogel rheology is measured by a stress-controlled rheometer (Malvern Kinexus Pro) with a 20 mm-diameter geometry. Sandpapers are glued on the top and the bottom plates of the rheometer to minimize the sliding of the hydrogel during the rotational measurements. Fully swollen hydrogels are cut into 20 mm circular pieces by a tissue puncher to match the geometry size. All the rheology measurements are done at 37°C with a humidity cover. The single frequency tests are conducted with 1 Hz frequency and 1% shear strain. The strain-dependent tests are conducted with 1 Hz frequency, and the frequency-dependent tests are conducted with 1% shear strain.

### Cell culture

Bovine aortic endothelial cells (BAEC), human mesenchymal stem cells (hMSC), and mouse embryonic fibroblast (mEF) cells are cultured in DMEM with 10% fetal bovine serum, 50 μg/ml penicillin-streptomycin, and MEM nonessential amino acids (Thermo Fisher Scientific). Cells are maintained in cell culture incubator with 95% humidity at 37°C with 5% CO_2_. During experiments, cell culture medium is changed every two days. Cell growth rate is represented by the cell densities at different days, which are determined by manually counting the cell numbers on a given imaging area.

To harvest whole cell proteins, BAECs or hMSCs grown on the large hydrogels are first detached from the hydrogel by 0.025% trypsin and 2 mM ethylenediaminetetraacetic acid (EDTA, Sigma-Aldrich). Cells are centrifuged, washed with PBS once and centrifuged again to remove the excess cell culture medium and trypsin. Each cell pallet is then lysed by cell lysis buffer at 95°C containing 1% sodium dodecyl sulphate (SDS), 10 mM Tris, and 1 mM EDTA at pH 8. Cell lysis solutions are sonicated and heated at 95°C for 20 min. The whole-cell protein concentrations are measured by the BCA protein assay (Pierce, Thermo Fisher Scientific). For SDS-polyacrylamide gel electrophoresis (SDS-PAGE), 20 μg whole-cell proteins are loaded onto the well of the gel and are stained with Coomassie blue. For Western blot analysis, 17 μg whole-cell proteins are loaded onto each well. The following antibodies are used: rabbit anti-filamin A (Novus Biologicals, Centennial, CO), mouse anti-gelsolin (Sigma-Aldrich), mouse anti-lamin A/C and rabbit anti-vimentin (Cell Signaling Technology, Danvers, MA), and mouse anti-β-actin (Abcam, Waltham, MA).

### Immunofluorescence staining

Fibroblast cells on hydrogels are fixed with 4% paraformaldehyde for 15 min, followed by being permeabilize with 0.2% triton-X for 10 min. Cells are then incubated with DyLigh 594 Phalloidin (1:500, Cell Signaling Technology) and DAPI (40 μg/ml, Sigma-Aldrich) for 20 min. Samples are fluorescently imaged by an upright fluorescence microscope (Nikon, 10×/0.45 lens).

### Traction force microscopy

For cell traction measurement, hydrogel pre-gel solution is mixed with 0.2 μm fluorescent beads (carboxylate-modified, Invitrogen, Thermo Fisher Scientific) with a concentration of 0.24% v/v. To quantify the hydrogel deformation caused by the contraction of a cell, two sequential fluorescence images for the fluorescent beads are taken, with one showing the cell spreading and contracting the gel and one after removing the cell from the hydrogel. The hydrogel deformation field is then calculated by particle image velocimetry, and the cell traction force is reconstructed by solving the Boussinesq solution, both of which are conducted by open-source plugins in NIH ImageJ ^[21]^. Average traction forces are determined as the summation of the amplitude of the local traction forces beneath the cell projected region divided by the cell projected area.

### Statistical analysis

Statistical significance is analyzed using unpaired, two-samples Student’s t-test.

